# Improved gene tree inference from removing alignment errors both from focal genes and when training substitution models

**DOI:** 10.64898/2025.12.01.691663

**Authors:** Andrew L. Wheeler, Chiragdeep Chatur, Peter W Goodman, Robert C Edgar, Gavin A Huttley, Joanna Masel

**Affiliations:** Genetics Graduate Interdisciplinary Program, University of Arizona, Tucson, Arizona 85721, USA; Department of Computer Science, University of Arizona, Tucson, Arizona 85721, USA; Statistics & Data Science, University of Arizona, Tucson, Arizona 85721, USA; Independent researcher, Corte Madera, California, USA; Research School of Biology, The Australian National University; Department of Ecology and Evolutionary Biology, University of Arizona, Tucson, Arizona 85721, USA

**Author notes:** **Corresponding author:** Joanna Masel.

**Keywords:** phylogenetic methods, phylogenomics, detecting selection, substitution matrix, alignment confidence, homology

## Abstract

Multiple Sequence Alignment (MSA) is a key step in phylogenetic analysis, and is prone to error. Unfortunately, algorithms that remove likely alignment errors from MSAs sometimes also remove informative residues, making phylogenetic tree inference worse. Here we present a novel MSA cleaning algorithm based on consensus between MSAs using a range of Hidden Markov Models and guide trees, named CLOAK (<u>CL</u>eaning <u>O</u>n the basis of <u>A</u>lignment C(<u>K</u>)onsensus). CLOAK is a gentle filter, with a low false positive rate for removal from MSAs according to the BALiBASE benchmarks, while still removing a significant fraction of likely alignment errors. Gentle vs. stringent MSA filtering methods are appropriate for different tasks. We assess methods based on their ability to bring the gene trees of single copy orthologs closer to the accepted species tree. Amino acid substitution models trained on filtered MSAs improve gene tree inference, with stricter filtering methods providing the biggest model improvements. In contrast, it is gentler filtering of single gene MSAs that provides additional improvements to gene tree inference, with CLOAK performing best.

## Introduction

Multiple sequence alignment (MSA) is an important step in processing sequence data, prior to performing evolutionary inference. MSA attempts to identify homologous sites (Morrison, et al. 2015), allowing the use of models of the evolutionary history of substitutions at each site. The alignment of non-homologous sites will propagate to many types of downstream analysis that assume homology, including inference of phylogenetic trees and of selection (Wong, et al. 2008). Estimates of positive selection are particularly sensitive to the alignment method used, and alignment errors can inflate false positive rates for positive selection (Schneider, et al. 2009; Fletcher and Yang 2010; Jordan and Goldman 2012). The inference of phylogenetic trees can also depend on the alignment method used (Wong, et al. 2008). Tree accuracy can be improved by co-inferring the MSA with the phylogenetic tree, thus appropriately accounting for MSA uncertainty (Liu, et al. 2009; Shim and Larget 2018). Unfortunately, this increases computational expense, and does not scale well to larger datasets. Another option is to abandon likelihood-based approaches in favor of distance-based phylogenetics whose distance metric comes from a Hidden Markov Model rather than a fixed MSA (Bogusz and Whelan 2017). However, the two-step approach of first generating an MSA and then taking it as fact while performing evolutionary analysis remains standard practice.

Given that MSA errors are inevitable (Landan and Graur 2009), an attractive approach is to filter out residues that are likely to be misaligned, prior to downstream analysis. Unfortunately, alignment filtering can counterproductively lead to worse performance on phylogenetic inference (Tan, et al. 2015; Portik and Wiens 2021) and selection detection (Spielman, et al. 2014). For a single gene tree (rather than a species tree based on a large, concatenated alignment), inadvertently removing even a small number of correctly aligned sites further reduces the availability of already scarce information.

MSAs are also used as training data for empirical substitution models, which specify the rates with which each nucleotide, amino acid, or codon evolves into each alternative state. Misaligned amino acids might introduce systematic errors into substitution models, e.g. inflating the inferred rate of unlikely substitutions. Substitution rates might also become inflated for amino acid pairs that are most likely to be aligned as the least bad option within a highly diverged sequence for which alignment confidence is low. Indeed, alignment uncertainty has been found to affect fit of substitution model (Spielman and Miraglia 2021). To the best of our knowledge, the impact of filtering errors out of training MSAs for substitution models has not been assessed.

Substitution models enable the use of likelihood-based methods to infer trees (Felsenstein 1981), and are also used in some distance-based methods (Bogusz and Whelan 2017). Substitution model choice thus impacts phylogenetic inference. While common practice is to use a tool such as ModelFinder (Kalyaanamoorthy, et al. 2017) to choose among existing substitution models, the QMaker program within IQ-Tree recently made it straightforward to train custom substitution models (Minh, et al. 2021). Here, we test whether removing alignment errors from the MSA training data input into QMaker improves subsequent gene tree inference.

The earliest approaches for MSA filtering focused on removing entire columns from an alignment. Gblocks removes poorly conserved columns that are found near gaps or are contiguous with other poorly conserved sites (Castresana 2000). TrimAl eliminates columns that fall below a score threshold, using several possible criteria such as number of gaps and residue similarity (Capella-Gutiérrez, et al. 2009). Other methods score how phylogenetically informative a column is (Dress, et al. 2008), randomness or entropy (Criscuolo and Gribaldo 2010; Kück, et al. 2010), or alignment uncertainty (Penn, et al. 2010).

Because these methods use an “all or nothing” approach for each column within an alignment, they will be too strict in a column where a subset of the sequences in a MSA are well aligned, but other sequences are poorly aligned. Deleting whole columns at such sites can result in removing information about closely related species, which is disproportionately found within rapidly evolving regions. These include intrinsically disordered regions, which tolerate a high rate of indels (de la Chaux, et al. 2007; Khan, et al. 2015). Tasks such as phylogenetic inference that are sensitive to loss of information are likely to benefit from a gentler approach that removes clear errors while retaining as many correctly aligned sites as possible.

Other methods identify stretches of likely non-homologous characters within each sequence. PREQUAL is a popular “pre-alignment” option that uses a Hidden Markov Model to identify stretches of sequence that are unlikely to be homologous to any other sequence in the dataset (Whelan, et al. 2018). This approach is focused on removing frameshift errors from sequencing, assembly errors creating mosaics, and/or unique insertions, rather than on removing alignment errors.

Divvier (Ali, et al. 2019) and HmmCleaner (Di Franco, et al. 2019) attempt to identify incorrectly aligned sequences by using probabilistic pair Hidden Markov Models to discern pairwise homology, rather than whole-column homology. These programs are influenced by earlier tools such as Zorro (Wu, et al. 2012), and PSAR (Kim and Ma 2014). Divvier offers the option to retain additional phylogenetic information by “divvying,” meaning splitting some columns into two or more internally well aligned subsets; we refer to this as Divvier-d. An alternative setting in Divvier uses an approach closer to HmmCleaner, retaining only the largest subset; we refer to this as Divvier-pf. TAPER takes a different approach, searching for sliding windows of sequence that the alignment suggests are unusually divergent relative to what is expected for that species (Zhang, et al. 2021). An important question is whether even these methods remove too much true phylogenetic signal, thus worsening gene tree inference.

Consensus among alternative alignments is another potentially useful source of information. Alignment consensus tools such as MergeAlign (Collingridge and Kelly 2012), and M-Coffee (Wallace, et al. 2006) do not filter out any residues, but generate one consensus alignment from an ensemble.

Alternatively, MSA uncertainty can be propagated, and consensus taken later, e.g. among trees. This might be an effective approach to filter out random errors in an alignment, but if there are also significant systematic errors across the variant MSAs, these would require filtering. This approach will also substantially increase computational cost.

Here we propose identifying likely alignment errors based on departure from consensus within an ensemble of alternative alignments. This is implicit in Divvier’s and HmmCleaner’s use of HMM confidence. We also include another source of alignment uncertainty: that in the “guide tree” that specifies the order in which sequences are added during MSA construction. The guide tree is often a neighbor joining or UPGMA tree that groups together more similar sequences (Feng and Doolittle 1987). Variation in which guide tree is used can contribute substantial variability in the resulting MSA (Zhan, et al. 2015). The alignment cleaning tool GUIDANCE2 generates a set of perturbed alignments by leveraging uncertainty in the guide tree. It combines this with simple penalties for alignment gaps, rather than a full HMM approach, to compute reliability scores. This enables unreliable alignments to be filtered out, with the option of filtering only parts of columns (Sela, et al. 2015).

Here we develop a new consensus-based approach to alignment filtering. We look for consensus among alternative MSAs corresponding to perturbations of both the HMM and the guide tree, using an ensemble of alignments generated by Muscle5 (Edgar 2022). Pairings that appear inconsistently across these alignments indicate uncertainty and hence suggest unreliable alignment. In this light, we introduce a novel algorithm, named <u>CL</u>eaning <u>O</u>n the basis of <u>A</u>lignment C(<u>K</u>)onsensus (CLOAK). For maximal information retention, we adopt the practice of divvying (Ali, et al. 2019).

We assess the value of filtering, both in the construction of substitution models and in the inference of a gene tree, in terms of whether the resulting gene tree of a single copy ortholog is closer to the ground truth we expect from a known species tree. We use a mammalian set of single copy orthologs (Allio, et al. 2024), across a set of species diverged enough to have little incomplete lineage sorting. We are therefore able to test how MSA cleaning affects the output of interest, namely the accuracy of the gene tree.

Best practices are with respect to alignment filtering might depend on the application. We expect substitution model inference, which uses a large set of training MSAs, to tolerate aggressive filtering approaches that also remove a significant number of true homologies, in order to maximize the accuracy of residues that remain. For inferring gene trees, losing any well-aligned residues could harm the already limited information available for inference, requiring a lighter touch to filtering. Here we identify best practices, judged according to bringing inferred ortholog trees closer to the known species tree.

## Results

### CLOAK removes fewer correctly aligned residues than other alignment filtering tools

One way to assess an MSA is to compare it to a benchmark that putatively represents the ground truth alignment. A good MSA, or a well-filtered MSA, will maximize the number of pairings that match the ground truth, while minimizing the number of pairings that do not. We use BALiBASE, whose alignments are based on protein structure, as our benchmark (Thompson, et al. 2005). We evaluate alignments, both before and after filtering, using a precision-recall curve. Here precision on the x-axis is equal to 1 minus the false discovery rate in which amino acids are aligned in a manner not found in BALiBASE, and recall on the y-axis represents the rate that (presumed correct) BALiBASE pairings are recovered (Figure 1). We confirm previous results indicating that Muscle5 outperforms MAFFT and Clustal Omega, with superiority indicated by both higher precision and higher recall (Edgar 2022). Therefore, we use Muscle5 alignments as our starting point, prior to filtering.

**Figure 1.**
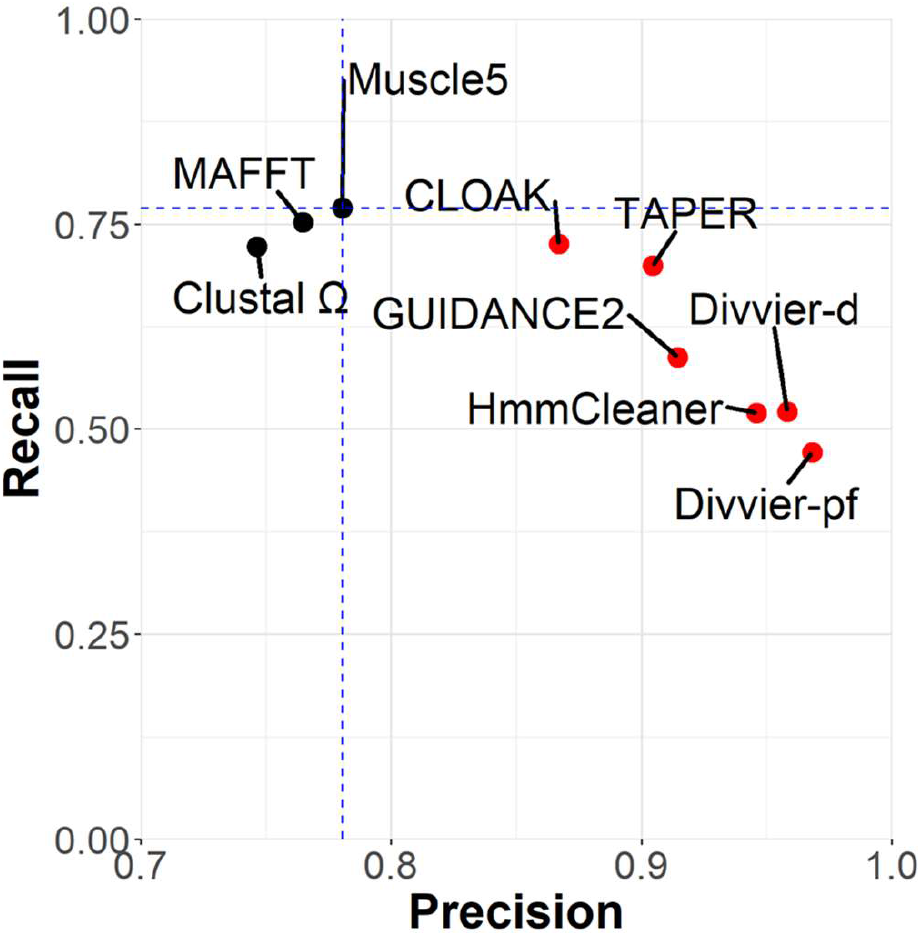
CLOAK removes a significant fraction of false pairings, with minimal loss of true information. The recall, or true positive rate, is the fraction of aligned pairs in the core region of the BAliBase reference alignment that are recovered. The precision, equal to 1 minus the false discovery rate, is the number of aligned pairs matching the BAliBase reference divided by the total number of aligned amino acid pairs in the BAliBASE core alignment. Different alignment programs are shown as black points, and different filtering methods are shown in red. Each of the filtering methods uses the Muscle5 alignment as its starting point, with distance from the blue horizontal and vertical dashed lines highlighting the extent to which true and false positives are filtered.

HmmCleaner combines lower precision with similar recall to Divvier – we therefore do not recommend HmmCleaner. Using GUIDANCE2 to remove residues with a confidence score <0.9 (see methods) is similarly inferior to TAPER, with an only slightly higher precision at the cost of substantially worse recall. CLOAK, TAPER, Divvier-d (with divvying), and Divvier-pf (with partial filtering) all fall along a reasonable curve, where the choice of filtering program among these options may vary among applications, depending upon what balance of false positives (precision) and false negatives (recall) each application calls for.

Divvier-pf removes the most error at the cost of losing more correct homologies, making it the best option for applications that are highly sensitive to error, but which have abundant available data such that data loss is tolerable. We expect substitution model inference to fall into this category. CLOAK has the gentlest touch, removing less error but removing almost no pairings scored as correct by BALiBASE; CLOAK may be more suited for applications with highly limited available data, such as inferring single gene phylogenies.

### Alignment filtering lowers inferred rates of less mutationally accessible substitutions

In the standard genetic code, some amino acid substitutions require only a single nucleotide substitution, while others require 2 or 3 point mutations (in the absence of insertions or deletions, which occur at lower rates (Chen, et al. 2009)). To avoid the confounding influence of amino acid frequencies, we quantify the accessibility of substitutions corresponding to different mutational accessibilities according to “exchangeabilities”. The flux of events in which amino acid A evolves into amino acid B is equal to the frequency of A times the rate *k*_*AB*_, and B evolves into A with flux *Bk*_*BA*_. An exchangeability is defined as 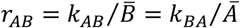, such that at equilibrium 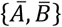, the fluxes in both directions are both equal to 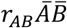. Exchangeabilities tend to be lower between amino acid pairs that are less mutationally accessible (Supplementary Figure 1). However, we expect MSA errors to introduce systematic errors that make a proportionately larger contribution to mutationally inaccessible and hence rare substitutions. Indeed, as predicted, filtering MSAs preferentially lowers the inferred exchangeabilities of less accessible substitutions (Figure 2). Again, as predicted, this effect is stronger with stricter filters such as Divvier-pf (Figure 2A, Spearman’s ρ =-0.54) than with a gentler filter such as CLOAK (Figure 2B, Spearman’s ρ =-0.41). The intermediate strength filters Taper and Divvier-d also follow the expected dose-response across the range of filtering strengths, with Spearman’s ρ of -0.46 and -0.50, respectively.

**Figure 2.**
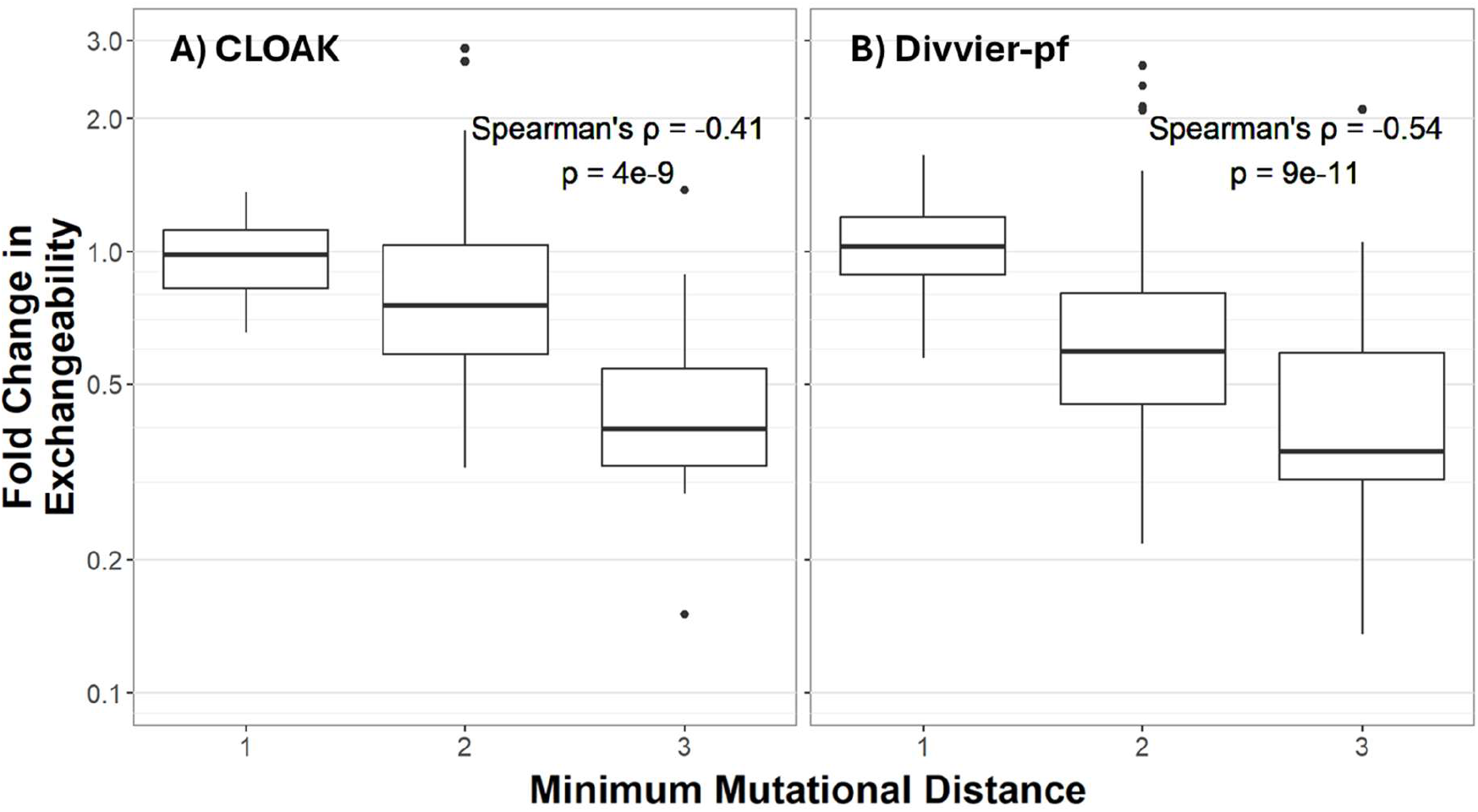
Rates of implausible substitution drop more after filtering alignments with the stricter Divvier-pf than with the gentler CLOAK. Implausible substitutions are those that require a minimum of 2 or 3 nucleotide point mutations. Exchangeabilities come from substitution models trained on a filtered vs. unfiltered alignments of mammalian orthologs (see Methods). Boxes show 25% and 75% quartiles, whiskers show 95% confidence intervals, and the central tendency line shows the median.

### Filtering Improves Tree Inference

The most important test of alignment filtering is not whether it brings alignments closer to a still-uncertain benchmark, but whether it improves inference, e.g. of phylogenetic trees. We use the accepted mammalian species tree (Upham, et al. 2019) as ground truth to assess the accuracy of gene trees of mammalian single copy orthologs, which are expected to follow the species tree.

When using substitution models trained on filtered data, gene trees are more similar to the corresponding species tree (Figure 3A, D, and G; alignments to the right of the vertical black line are improved by filtering). We get more statistical power using the Lin-Rajan-Moret distance (Lin, et al. 2012) (Figure 3 first column; see p-values in table 1) than from using the proportion of gene tree quartets (Estabrook, et al. 1985; Day 1986) that agree with the species tree topology (Mutti, et al. 2025) (Figure 3 middle column), or the number of duplication, loss, and transfer events inferred by gene tree-species tree reconciliation (Figure 3 right column). The Lin-Rajan-Moret distance is a weighted extension of the more commonly used Robinson-Foulds distance metric (Robinson and Foulds 1981) that offers superior discrimination and robustness to small changes, such as moving a single leaf (Lin, et al. 2012).

**Table 1.**
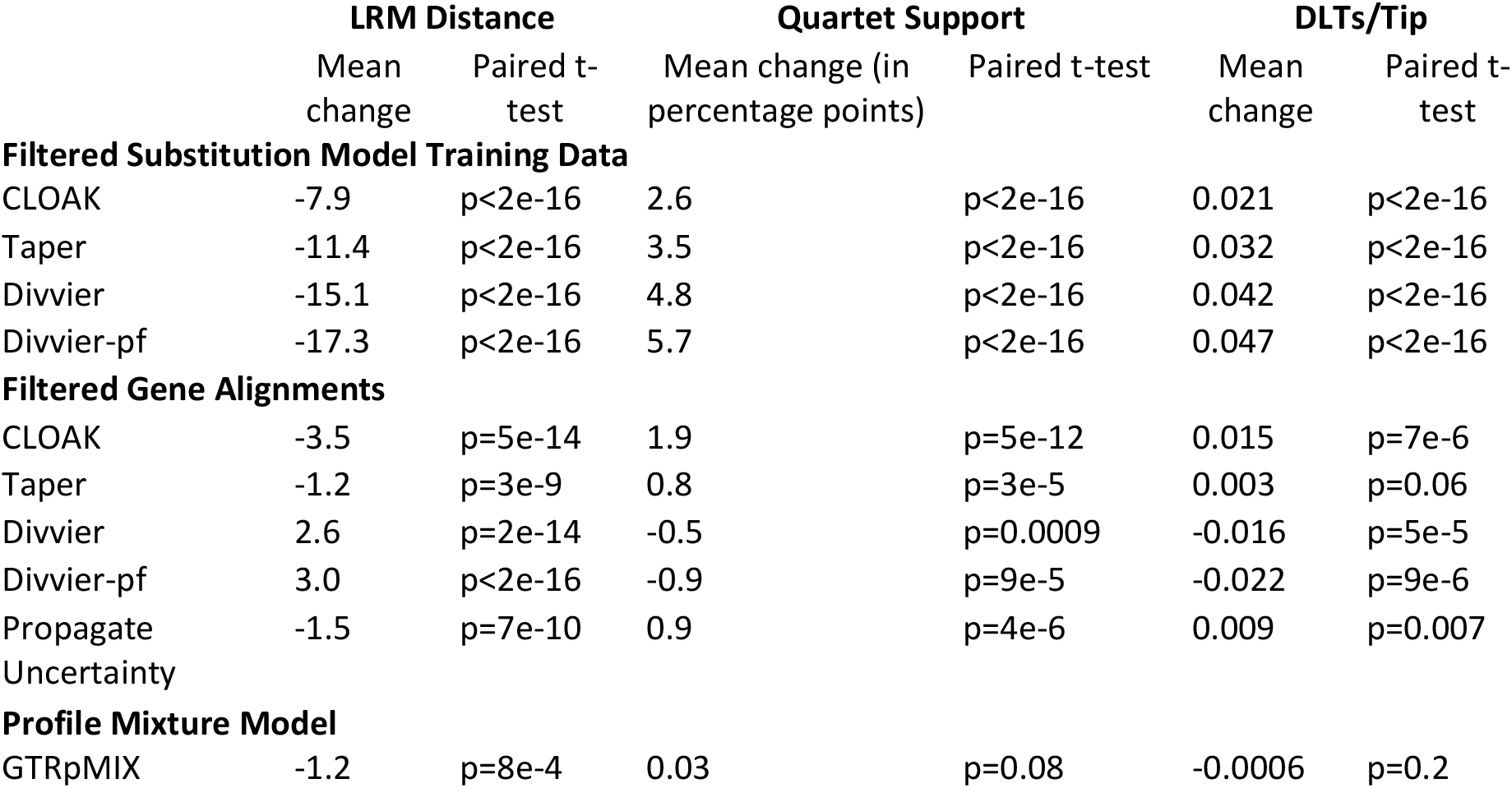
Effect sizes and significance of all comparisons in Figure 3.

**Figure 3.**
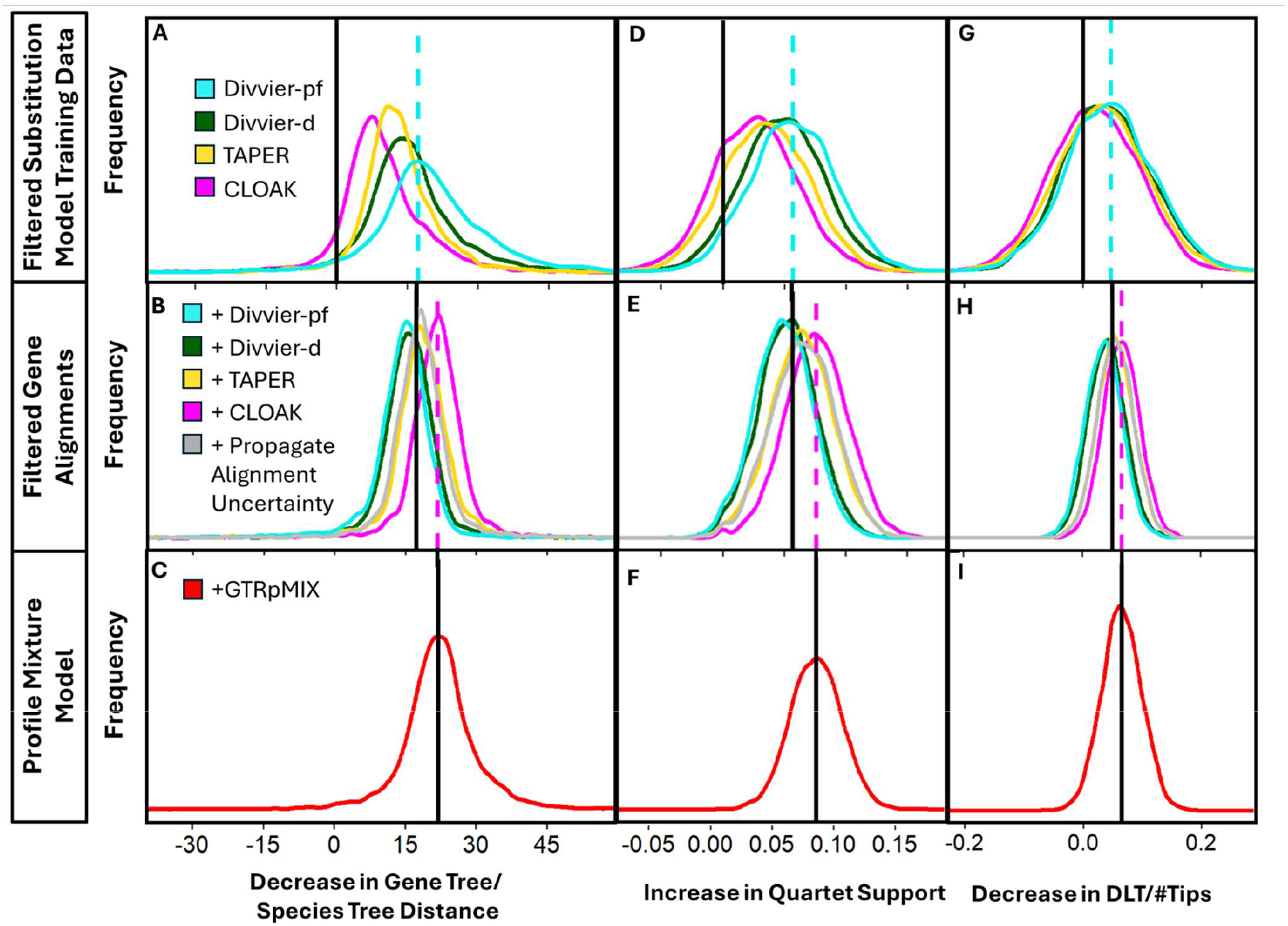
Filtering of MSAs improves tree inference. We measure the similarity of each mammalian single copy ortholog tree to the mammalian species tree. A rightward shift indicates increased similarity following filtering. The figure shows the distribution of pairwise differences calculated for each gene. The first column (panels A,B, and C) shows decreases in the Lin-Rajan-Moret distance (Lin, et al. 2012). The second column (panels D, E, and F) shows the increase in the proportion of gene tree quartets that agree with the species tree topology. The third column (panels F, H, and I) shows decreases in duplication, loss, and transfer (DLT) events inferred by reconciling each gene tree with the species tree. Baseline gene trees were inferred using unfiltered alignments for individual genes, as well as a substitution model trained on unfiltered data. All effect sizes and p-values can be found in table 1; the Lin-Rajan-Moret distance produced the most statistical power. By all three metrics, the more stringent the filtering on substitution model training data, the greater the gain in gene tree accuracy (top row, panels A, D, and G). I.e., Divvier-pf produced the greatest accuracy gains, followed by Divvier-d, then TAPER, then CLOAK. Gentle filtering of single gene MSAs yields additional gene tree improvements (middle row, panels B, E, and H). Baseline trees in the middle row are the best case from the panel above, using unfiltered single gene alignments and a substitution model trained on Divvier-pf filtered data. Trees improved when filtering gene alignments with CLOAK and to a lesser extent TAPER, as well as when propagating alignment uncertainty to produce a consensus tree downstream. However, gene trees become worse following more stringent filtering with Divvier-d or Divvier-pf. Using the profile mixture model GTRpMIX (bottom row, panels C, F, and I), provides only marginal improvement in LRM distance, with no significant change in quartet support or duplication, losses, and transfers.

The best-performing substitution model comes from applying the strictest filter, Divvier-pf, to the substitution model training data (Figure 3, top row). Applying any of the four high-performing filtering methods (chosen above on the basis of BALiBASE) to substitution model training data leads to some improvement in gene trees.

Filtering the individual MSAs used for gene tree inference yielded additional improvement when using the gentler CLOAK and TAPER filters, but not when using the stricter Divvier-d or Divvier-pf (Figure 3B, E, and H; vertical black line indicates the use of Divvier-pf in generating substitution models, additional improvement is indicated by a further shift to the right). The gentlest filtering option, CLOAK, performed best (pink histogram), although the additional reduction in Lin-Rajan-Moret distance is only ∼20% of the magnitude of that already obtained from using Divvier-pf filtered data to train substitution models.

Propagating the uncertainty represented by the set of 16 Muscle5 alignments into 16 trees tended to slightly improve the resulting consensus gene tree (grey histogram in Figure 3B, E, and H), but the improvement is smaller than that obtained by using the same 16 alignments within CLOAK to filter the MSA. This suggests the existence of systematic error across all 16 MSAs, rather than merely random error.

Profile mixture models, which use a distribution of amino acid frequency vectors across sites, have been shown to perform better than the LG matrix (Banos, et al. 2024). We inferred a profile mixture model from our Divvier-pf filtered training MSA dataset, then refitted the profile mixture on the CLOAK filtered test set gene alignments. This procedure yielded only marginal improvement according to Lin-Rajan-Moret distance, and no significant change according to quartet support or number of duplications, losses, and transfers, relative to a comparable filtering pipeline without use of a profile mixture (Figure 3C, F, and I, red histogram vs vertical black line).

Figure 4 shows absolute tree quality with no filtering, and with our best-performing filters, whereas Figure 3 shows change in tree quality. The relatively small shifts in Figures 4a-4c illustrate that our statistical power in Figure 3 stems from its reliance of paired analyses – variation among genes is high. This is unsurprising, e.g. genes vary in length and hence information content.

**Figure 4:**
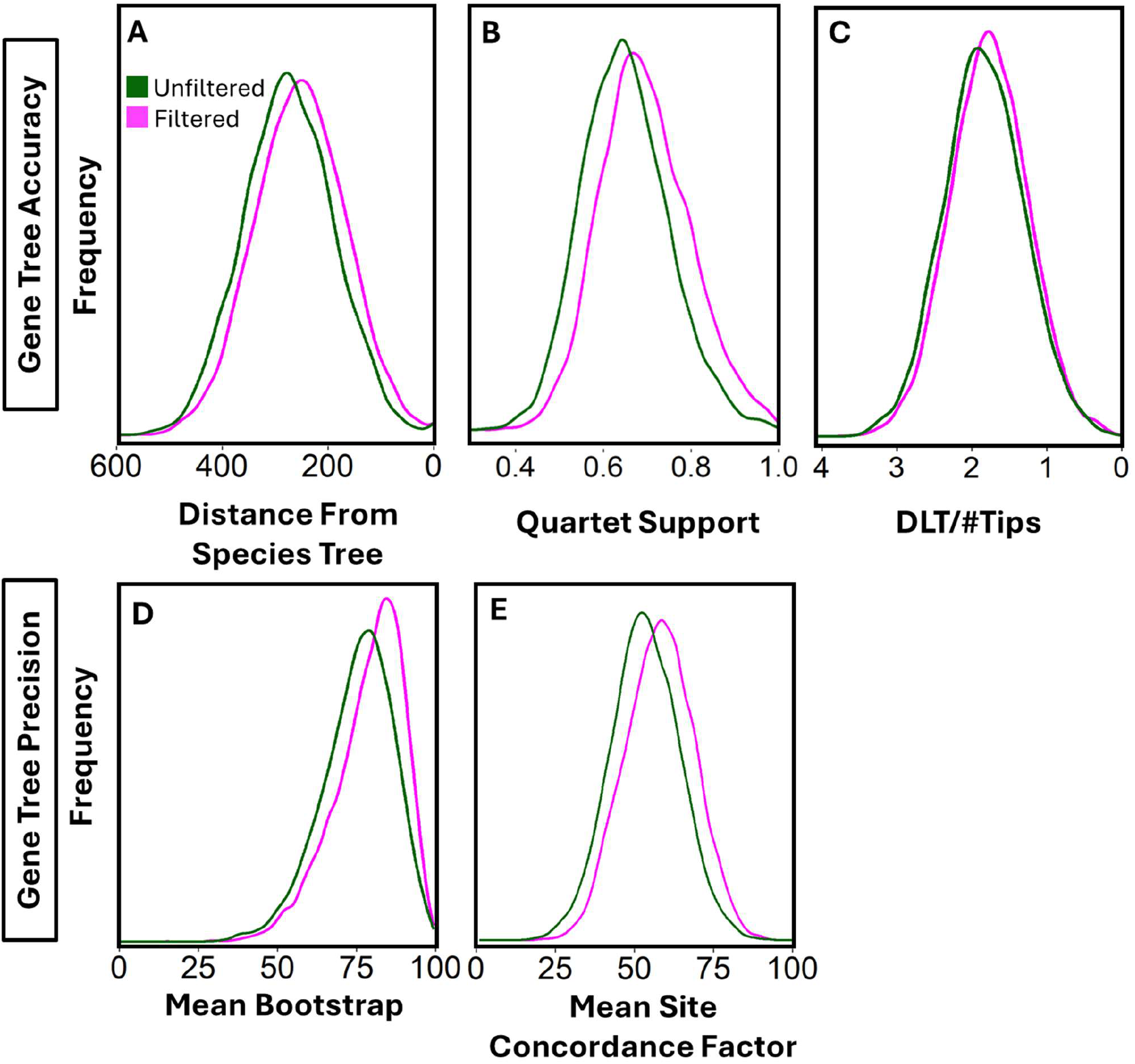
Absolute improvement in tree quality is modest relative to high variance in gene tree quality. Unfiltered gene trees correspond to the baseline case indicated by the vertical black line in Figure 3A/D/G, with filters for neither the gene alignments nor the substitution model training data. Filtered gene trees use gene alignments filtered with CLOAK, and a substitution model trained on data filtered with Divvier-pf (best practice pink curve in Figure 3B/E/H). Lin-Rajan-Moret distance is used in (A). Measures of tree confidence also show modest improvement (D: difference in means=3.4%, E: difference in means=4.6%) following the same filtering regime; all panels produce p<2e-16 in a paired t-test.

When ground truth in the form of a species tree is unavailable, metrics of tree confidence such as bootstrap confidence scores are sometimes used to assess tree quality. Mean bootstrap scores reflect genuine tree improvements in our analysis (Figure 4d). Another approach to quantify precision is to use site concordance factors, quantifying the fraction of informative sites in an alignment that support the tree topology at each node (Mo, et al. 2023). Bootstrap scores are inflated when there is genuine incongruence in a large dataset (Rokas and Carroll 2006); site concordance factors avoid this issue. Mean site concordance factors also reflect genuine tree improvements in our analysis (Figure 4e).

### Model fit does not predict model performance

A program such as ModelFinder is normally used to choose the best fitting substitution model under the presumption that this will generate the best tree (Kalyaanamoorthy, et al. 2017). However, if we do that with our CLOAK filtered gene alignments, the best-performing Q matrix (Divvier-pf filtered) is chosen less often (for 2405 out of 6769 genes or 36%) than the Q matrix trained on unfiltered data. This corresponds to the fact that CLOAK is a much gentler filter than Divvier-pf, and corresponding alignments are thus closer to unfiltered alignments.

## Discussion

The best approach to filtering MSAs depends on the application. We find that strictly filtering training data can significantly improve substitution models in a way that leads to more accurate phylogenetic tree inference. While previous studies have found that filtering the scarce MSA data used in gene tree inference is actually harmful (Tan, et al. 2015; Portik and Wiens 2021), we find that this is only true of strict filters, and that gentler filtering improves gene tree inference. We also present a new MSA filtering algorithm, CLOAK, based on consensus between alignments that perturb both HMMs and guide trees, that performs well as a gentle filter.

These recommendations from our work are downstream and distinct from early pipeline steps to remove sequences that may not be homologous at the whole-sequence level, or that are of poor sequence quality. A pre-alignment tool such as PREQUAL (Whelan, et al. 2018) can be helpful in such cases.

One concern when filtering alignments, especially when using the divvying function of CLOAK or Divvier-d, is that adding more gaps might make downstream inference worse than it would be if the column were simply omitted. Gaps are treated in most likelihood models as real amino acids or nucleotides whose identity is unknown, represented by an equilibrium frequency vector of possible amino acids (Felsenstein 1981). Gappier sequences will therefore tend to be inferred to be too similar to each other, and to sequences with typical amino acid usage. However, despite the fact that the divvying procedure of CLOAK introduces more gaps than many alternative approaches, it outperforms other methods for tree inference.

Our recommendations imply a claim about which masked alignment is “better” than another. What it means for an alignment to be better is not trivial (Morrison, et al. 2015). A common interpretation of an MSA is that characters in the same column are homologous, meaning they are descended from a shared ancestral nucleotide or amino acid. The alignment, including “-” characters, can represent point mutations, insertions, or deletions at that site.

However, for some evolutionary events, homology cannot be represented in MSA format. Following a tandem duplication, two characters in one sequence are homologous to one character in another. In an MSA, this would be represented as an insertion, with only one of the duplicates correctly shown as homologous to the original. Similarly, inversions can only be represented in an MSA as either a string of consecutive substitutions, or as a deletion and an insertion. In neither case does the MSA correctly identify the homology of the characters within the inversion. In amino acid or codon alignments, insertions and deletions at the nucleotide level might not match codon boundaries. For example, an out-of-frame deletion of 3 nucleotides creates a new codon formed from a hybrid of 2 other codons, and therefore an amino acid that is homologous to 2 ancestral amino acids (García Mesa, et al. 2024). Likelihood approaches will interpret the amino acid MSA as a point mutation plus an in-frame deletion.

With none of the plausible alignments able to capture the true evolutionary history at every site, we pragmatically judge alignment filtering methods based on their downstream ability to recover the correct gene tree. In this framework, the goal of filtering MSAs is not necessarily to make alignments “more homologous”, but to remove sites that impede accurate downstream inference.

Models trained on unfiltered data tend to be the best fit to unfiltered or lightly filtered sequence alignments, and are therefore chosen by popular model selection tools, even when this leads to less accurate trees. When alignment errors are amplified by an subsample–upsample approach (Sharma and Kumar 2022), then poor model choice might be particularly likely. One option we recommend using a stricter filter on alignments prior to using tools such as ModelFinder, separately from using a gentle filter after selecting the substitution model and before running tree inference. An alternative and increasingly available option is to make an *a priori* choice of taxon-specific exchangeability values that have been previously been trained on filtered data, set equilibrium amino acid frequencies to those in the current dataset, and use tools such as ModelFinder only to choose the model for rate variation among sites.

The optimal approach to filtering MSAs may be different for downstream applications other than gene tree inference. For example, tests for positive selection are very sensitive to alignment errors, resulting in a large number of false positives (Schneider, et al. 2009). Some studies have suggested that filtering with GUIDANCE can improve accuracy (Privman, et al. 2012) while others found little benefit (Jordan and Goldman 2012; Spielman, et al. 2014). However, stricter filtering with HmmCleaner was able to reduce the rate of false positives (Di Franco, et al. 2019). We therefore hypothesize that even stricter filtering using Divvier-pf would perform best for selection detection.

While we built CLOAK with amino acid alignments in mind, CLOAK can accept any fasta formatted alignment files, including nucleotide alignments, or even structural alignments generated by FoldMason using the 3Di alphabet (Gilchrist, et al. 2024). A similar consensus-based approach should be applicable to other alignment types, such as codon alignments, but CLOAK currently only accommodates fasta format. All that is needed is a set of alternative MSAs that capture uncertainty from an appropriate range of sources.

Most of the improvement we saw to gene tree accuracy came from using superior substitution models, rather than from filtering the gene MSA. This suggests that substitution model improvements are a promising avenue of research for improving phylogenetic inference. In the best case, a custom substitution model experimentally determined via deep mutational scanning of a specific protein dramatically outperform traditionally inferred models (Bloom 2014). Even without custom experimental data, bespoke substitution models trained for specific taxa outperform generic substitution models (Minh, et al. 2021; Dang, et al. 2022). A well-chosen alignment filter such as Divvier-pf can contribute to ensuring high quality data during the training of higher-performing substitution models.

## Methods

### CLeaning On the basis of Alignment C(K)onsensus (CLOAK)

CLOAK inputs a set of variant alignments of the same sequences and searches for consensus among them. We generate the input alignments for CLOAK using Muscle5, whose “-stratified” option conducts three perturbations of its core Hidden Markov Model (HMM) and three perturbations of its core guide tree, the combination of which produces 16 alternative alignments. Our algorithm retains each pair of amino acids in the same column if and only if they are found together (in the same column) in all 16 alternative alignments. If multiple subsets of amino acids within a column are found together across all alignments, but the whole column is not, the subsets are split into multiple columns that are each retained (divvying). Our method is shown in Figure 5.

**Figure 5.**
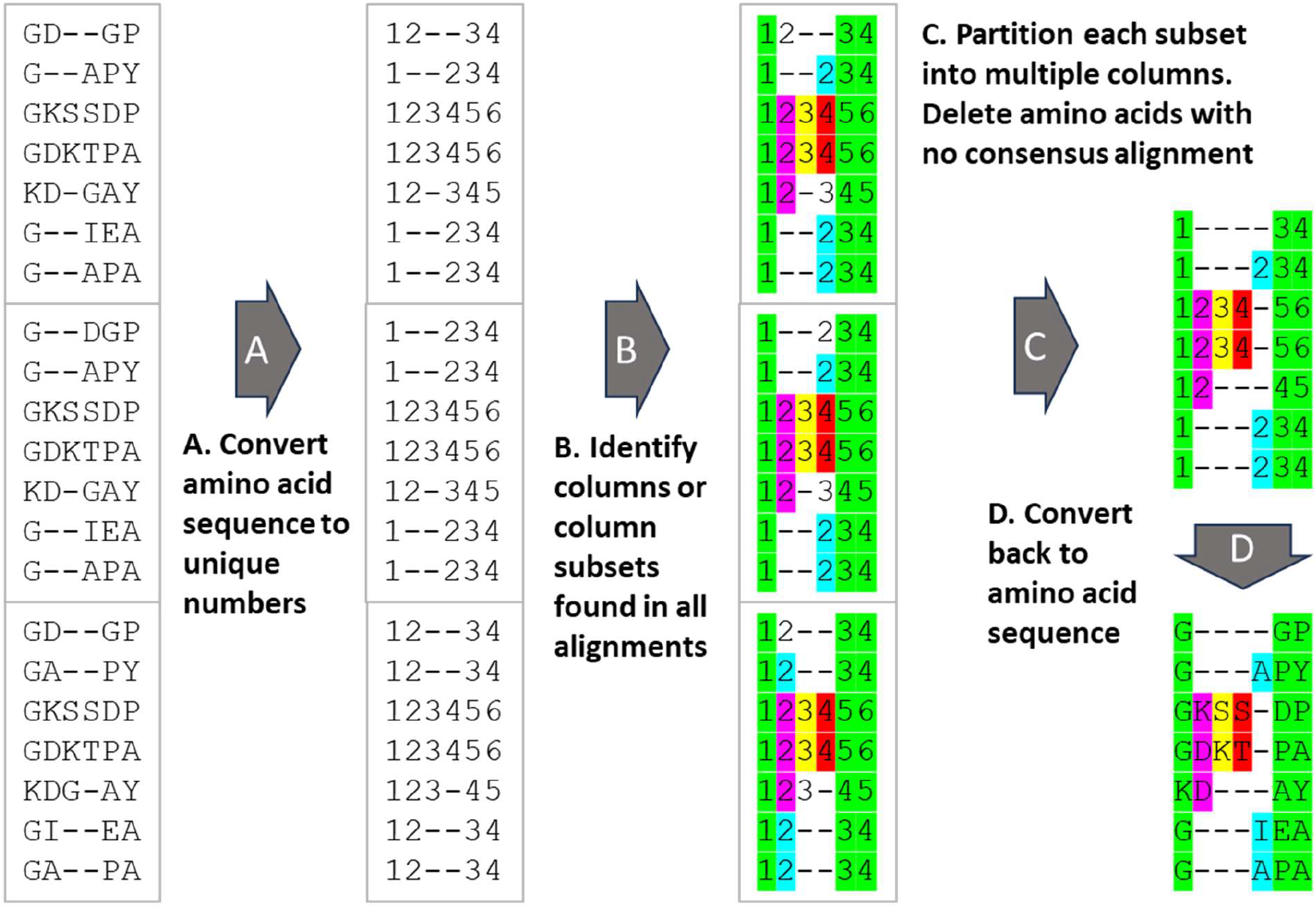
A schematic representation of our algorithm. Muscle5 produces 16 alternative alignments of the same amino acid sequences; we show only three for simplicity. To avoid confusion among amino acids of the same type, we first number each amino acid according to its position in the protein sequence. Our aim is to identify amino acid groupings shared in the same column within each alignment, albeit potentially in different columns in different alignments. We iterate over each column of the first alignment, proposing that all its amino acids be grouped together. We then check which column(s) those amino acids appear in each of the other alignments. When all amino acids in the column are not found together across all alignments, we divvy (Ali, et al. 2019) our proposed grouping into multiple, smaller proposed amino acid subgroupings, until a set of subgroupings is found across all alignments. We discard subgroupings of only a single amino acid because these contain no information. Other subgroupings are each given their own column. Finally, we convert numbers back into amino acids, to return a filtered, divvied alignment.

### Benchmarking alignments with BALIBase

We re-aligned the sequences in the BALiBASE version 3.0 dataset (Thompson, et al. 2005), including the N- and C-termini, using Muscle version 5.1 (Edgar 2022), MAFFT version 7.471, and Custal Ω version 1.2.4. We filtered each Muscle5 alignment using CLOAK, Divvier (using the -divvygap option, and with or without the partial filtering option) (Ali, et al. 2019), HmmCleaner version 0.243280 (Di Franco, et al. 2019), TAPER (Zhang, et al. 2021), and GUIDANCE version 2.01 (Sela, et al. 2015). GUIDANCE2 uses a set of variant alignments (by perturbing the guide tree), from which it calculates confidence scores for each residue. We used variant alignments from Muscle5, and masked all amino acids in the alignment with residue scores < 0.9. For all programs, default parameters were used except where specified.

We compared each filtered or unfiltered alignment to the structurally aligned region of BALiBASE (presumed to be ground truth), excluding the N- and C-terminal annotated regions, to quantify precision and recall (Figure 1). Our script for this was adapted from that used in Ali, et al. (2019) to assess Divvier (Simon Whelan, script provided via personal correspondence), which compares all pairwise homologies within a region shared by 2 alignments.

### Mammalian orthologs

To minimize the degree to which the true gene tree differs from the species tree (due to paralogy, incomplete lineage sorting, and horizontal gene transfer), we use a curated set of mammalian orthologs. We used a set of 6769 orthologous mammalian protein sequences across 190 species from the OrthoMaM database version 12 (Allio, et al. 2024), using only genes with at least one sequence in each of the 7 mammalian taxonomic groups annotated in the OrthoMaM database (euarchontes, glires, laurasiatheria, afrotheria, marsupialia, monotremata, and xenarthra). Starting with the unfiltered alignments from OrthoMaM, we re-aligned the sequences using Muscle5 (Edgar 2022). We used a subsample of 1000 of these loci to train amino acid substitution models, and the remaining 5769 loci as a testing set to infer single gene phylogenetic trees.

### Amino Acid Substitution Models

Amino acid substitution models describe a continuous time Markov process via a rate matrix **Q**, containing the rates at which each of the 20 amino acids are substituted for each other. In a time-reversible model, the fluxes between each member of an amino acid pair are equal in both directions. **Q** can then be conveniently expressed, using fewer parameters, as the product of a symmetric matrix of exchangeability rates (*r*_*ij*_ = *r*_*ji*_) and a vector of equilibrium amino acid frequencies (**π**), where *q*_*ij*_ = *r*_*ij*_π_*j*_ such that *q*_*ij*_π_*i*_ = *q*_*ji*_π_*j*_. QMaker takes equilibrium amino acid frequencies **π** to be equal to current amino acid frequencies (Minh, et al. 2021). The diagonal entries of **Q** (*q*_*i,i*_), are set such that each row sums to 0.

After inferring a substitution matrix with QMaker, we manually perform the normalization of **Q** that is normally done within IQ-Tree. We divide the matrix by a factor

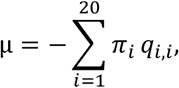

making the total number of substitutions per unit time equal to 1. We calculated normalized exchangeability values (for Figure 2 and supplementary Figure 1) from normalized **Q** matrices as:

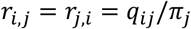

Because alignment filtering removes some amino acids more than others (e.g. preferentially removing amino acids that tend to be rapidly evolving), substitution models trained on filtered data will have equilibrium amino acid frequencies that do not match those of the original alignments. To avoid this problem, we generate **Q** by multiplying the exchangeability matrix trained on filtered MSAs by the equilibrium amino acid frequency vector calculated from the testing dataset. In some but not all cases, this testing dataset was also filtered, generally with a milder filter.

### Comparison of Gene Trees to Species Trees

Our mammalian species tree is a subset of the vertlife maximum clade credibility tree (Upham, et al. 2019), restricted to the 190 species in the OrthoMaM ortholog dataset (Allio, et al. 2024). The vertlife tree was constructed from a supermatrix of 31 gene alignments, drawing data from 4098 mammalian species.

We inferred gene trees from Muscle5 gene alignments of the testing subset of our mammalian ortholog sequences. Tree inference was performed using IQTree2 (Minh, et al. 2020), manually specifying which substitution matrix Q to use. For each gene, we selected a model for rate heterogeneity among sites using the ModelFinder program in IQTree2 (Kalyaanamoorthy, et al. 2017), which was usually a free rate model. To propagate alignment error, we inferred 16 alternative gene trees from an ensemble of 16 alternative Muscle5 alignments, and used the M-Coffee package of the T-Coffee distribution to infer a single consensus tree (Notredame, et al. 2000).

We calculated the Lin-Rajan-Moret distance (Lin, et al. 2012) between gene trees and the reference species tree using the *tree_distance* function in the cogent3 python library (Huttley, et al. 2025). The proportion of gene tree quartets that match the species tree was calculated with ASTRAL v.5.7.1 (Zhang, et al. 2018) using:

`java -jar astral.5.7.1.jar -i {gene treefile} -q {species treefile}`

The number of duplication, loss and transfer events in each gene tree was inferred with a parsimony approach implemented in Notung v.2.9.1 (Stolzer, et al. 2012), and normalized by the number of tips.

### Gene Tree Precision

We inferred bootstrap scores based on 1000 replicates per gene tree, using the ultrafast bootstrap application in IQTree2 (Hoang, et al. 2018). Site concordance factors are the proportion of informative sites in an alignment that support the tree topology at each node. We calculated site concordance factors for each gene tree using IQTree2 (Mo, et al. 2023), and took the mean across all branches in the tree.

## Code Availability

CLOAK is available both as an option within Muscle5 and as a stand-alone python script. Muscle5 can be found at https://www.drive5.com/muscle/, and CLOAK can be run within Muscle5 using the -cloak option. The stand-alone version of the CLOAK program, instructions for installing and running CLOAK alone and within Muscle5, and all scripts used for data analysis, can be found at https://github.com/phylowheeler/CLOAK.

## Author Contribution Statement

JM conceived the project, ALW and JM devised the general approach, CC refined and implemented the stand-alone Python version of CLOAK and produced most of Figure 1, PG integrated CLOAK into Muscle5 with oversight from RE, GH consulted on methods to assess tree accuracy, including suggesting the use of Lin-Rajan-Moret distance, ALW wrote the first draft with contributions from CC, RE, and JM, and ALW, and JM edited the final draft.

## Acknowledgements

We thank Simon Whelan for making available the code with which Divvier was assessed, which we modified for our own BAliBASE assessment. We thank Minh Bui, David Enard, Rob Lanfear, and Sawsan Wehbi for helpful discussions. We thank the National Science Foundation (DEB-2333243), the National Institutes for Health (T32 GM132008), and the University of Arizona Undergraduate Biology Research Program for funding.

## Supplemental Figures

**Supplementary Figure 1:**
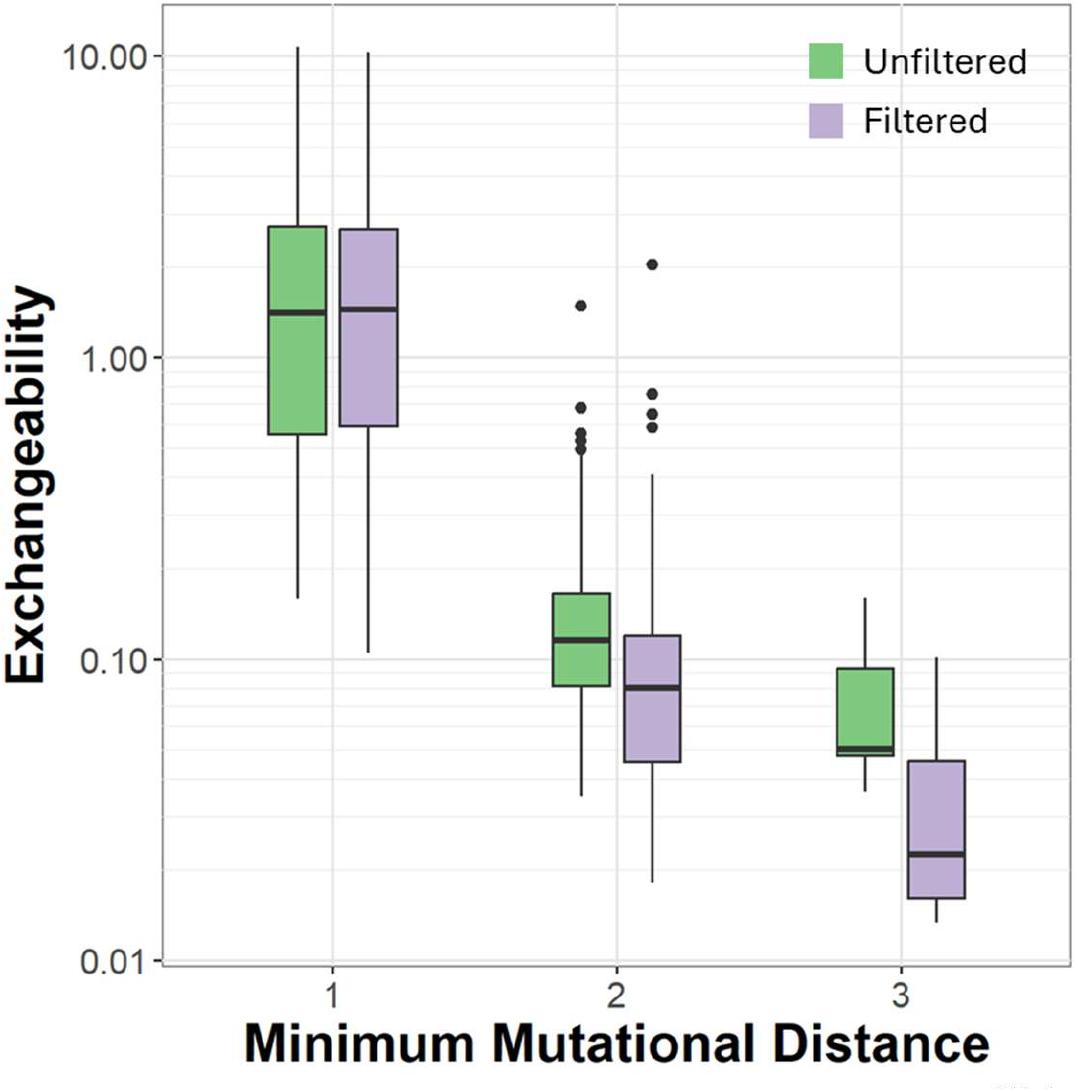
Exchangeabilities tend to be lower between amino acid pairs that are less mutationally accessible. The effects of filtering with Divvier-pf (green vs. purple) are smaller but still appreciable relative to exchangeability differences on the basis of minimum mutational distance. Minimum mutational distance indicates the minimum number of nucleotide point mutations required to achieve a given amino acid substitution. Exchangeabilities come from substitution models trained on a filtered vs. unfiltered alignments of mammalian orthologs (see Methods). Boxes show 25% and 75% quartiles, whiskers show 95% confidence intervals, and the central tendency line shows the mean.

## References Cited

Ali RH, Bogusz M, Whelan S. 2019. Identifying Clusters of High Confidence Homologies in Multiple Sequence Alignments. Mol Biol Evol 36:2340–2351.

Allio R, Delsuc F, Belkhir K, Douzery EJP, Ranwez V, Scornavacca C. 2024. OrthoMaM v12: a database of curated single-copy ortholog alignments and trees to study mammalian evolutionary genomics. Nucleic Acids Research 52:D529–D535.

Banos H, Wong TKF, Daneau J, Susko E, Minh BQ, Lanfear R, Brown MW, Eme L, Roger AJ. 2024. GTRpmix: A Linked General Time-Reversible Model for Profile Mixture Models. Molecular Biology and Evolution 41:msae174.

Bloom JD. 2014. An experimentally determined evolutionary model dramatically improves phylogenetic fit. Mol Biol Evol 31:1956–1978.

Bogusz M, Whelan S. 2017. Phylogenetic Tree Estimation With and Without Alignment: New Distance Methods and Benchmarking. Systematic Biology 66:218–231.

Capella-Gutiérrez S, Silla-Martínez JM, Gabaldón T. 2009. trimAl: a tool for automated alignment trimming in large-scale phylogenetic analyses. Bioinformatics 25:1972–1973.

Castresana J. 2000. Selection of Conserved Blocks from Multiple Alignments for Their Use in Phylogenetic Analysis. Molecular Biology and Evolution 17:540–552.

Chen J-Q, Wu Y, Yang H, Bergelson J, Kreitman M, Tian D. 2009. Variation in the Ratio of Nucleotide Substitution and Indel Rates across Genomes in Mammals and Bacteria. Molecular Biology and Evolution 26:1523–1531.

Collingridge PW, Kelly S. 2012. MergeAlign: improving multiple sequence alignment performance by dynamic reconstruction of consensus multiple sequence alignments. BMC Bioinformatics 13:117.

Criscuolo A, Gribaldo S. 2010. BMGE (Block Mapping and Gathering with Entropy): a new software for selection of phylogenetic informative regions from multiple sequence alignments. BMC Evol Biol 10:210.

Dang CC, Minh BQ, McShea H, Masel J, James JE, Vinh LS, Lanfear R. 2022. nQMaker: Estimating Time Nonreversible Amino Acid Substitution Models. Systematic Biology 71:1110–1123.

Day WHE. 1986. Analysis of Quartet Dissimilarity Measures Between Undirected Phylogenetic Trees. Systematic Biology 35:325–333.

de la Chaux N, Messer PW, Arndt PF. 2007. DNA indels in coding regions reveal selective constraints on protein evolution in the human lineage. BMC Evolutionary Biology 7:191.

Di Franco A, Poujol R, Baurain D, Philippe H. 2019. Evaluating the usefulness of alignment filtering methods to reduce the impact of errors on evolutionary inferences. BMC Evolutionary Biology 19:21.

Dress AW, Flamm C, Fritzsch G, Grünewald S, Kruspe M, Prohaska SJ, Stadler PF. 2008. Noisy: identification of problematic columns in multiple sequence alignments. Algorithms Mol Biol 3:7.

Edgar RC. 2022. Muscle5: High-accuracy alignment ensembles enable unbiased assessments of sequence homology and phylogeny. Nature Communications 13:6968.

Estabrook GF, McMorris FR, Meacham CA. 1985. Comparison of Undirected Phylogenetic Trees Based on Subtrees of Four Evolutionary Units. Systematic Zoology 34:193–200.

Felsenstein J. 1981. Evolutionary trees from DNA sequences: A maximum likelihood approach. Journal of Molecular Evolution 17:368–376.

Feng D-F, Doolittle RF. 1987. Progressive sequence alignment as a prerequisitetto correct phylogenetic trees. Journal of Molecular Evolution 25:351–360.

Fletcher W, Yang Z. 2010. The Effect of Insertions, Deletions, and Alignment Errors on the Branch-Site Test of Positive Selection. Molecular Biology and Evolution 27:2257–2267.

García Mesa JJ, Zhu Z, Cartwright RA. 2024. COATi: Statistical Pairwise Alignment of Protein-Coding Sequences. Molecular Biology and Evolution 41:msae117.

Gilchrist CLM, Mirdita M, Steinegger M. 2024. Multiple Protein Structure Alignment at Scale with FoldMason. bioRxiv:2024.2008.2001.606130.

Hoang DT, Chernomor O, von Haeseler A, Minh BQ, Vinh LS. 2018. UFBoot2: Improving the Ultrafast Bootstrap Approximation. Molecular Biology and Evolution 35:518–522.

Huttley G, Caley K, Fotovat N, Ma SK-W, Koh M, Morris R, McArthur R, McDonald D, Jaya F, Maxwell P, et al. 2025. Cogent3: Making Sense of Sequence. In: Zenodo doi: 10.5281/zenodo.16519079.

Jordan G, Goldman N. 2012. The effects of alignment error and alignment filtering on the sitewise detection of positive selection. Mol Biol Evol 29:1125–1139.

Kalyaanamoorthy S, Minh BQ, Wong TKF, von Haeseler A, Jermiin LS. 2017. ModelFinder: fast model selection for accurate phylogenetic estimates. Nat Methods 14:587–589.

Khan T, Douglas GM, Patel P, Nguyen Ba AN, Moses AM. 2015. Polymorphism Analysis Reveals Reduced Negative Selection and Elevated Rate of Insertions and Deletions in Intrinsically Disordered Protein Regions. Genome Biology and Evolution 7:1815–1826.

Kim J, Ma J. 2014. PSAR-align: improving multiple sequence alignment using probabilistic sampling. Bioinformatics 30:1010–1012.

Kück P, Meusemann K, Dambach J, Thormann B, von Reumont BM, Wägele JW, Misof B. 2010. Parametric and non-parametric masking of randomness in sequence alignments can be improved and leads to better resolved trees. Front Zool 7:10.

Landan G, Graur D. 2009. Characterization of pairwise and multiple sequence alignment errors. Gene 441:141–147.

Lin Y, Rajan V, Moret BM. 2012. A metric for phylogenetic trees based on matching. IEEE/ACM Trans Comput Biol Bioinform 9:1014–1022.

Liu K, Raghavan S, Nelesen S, Linder CR, Warnow T. 2009. Rapid and Accurate Large-Scale Coestimation of Sequence Alignments and Phylogenetic Trees. Science 324:1561–1564.

Minh BQ, Dang CC, Vinh LS, Lanfear R. 2021. QMaker: Fast and Accurate Method to Estimate Empirical Models of Protein Evolution. Systematic Biology 70:1046–1060.

Minh BQ, Schmidt HA, Chernomor O, Schrempf D, Woodhams MD, Von Haeseler A, Lanfear R. 2020. IQ-TREE 2: New Models and Efficient Methods for Phylogenetic Inference in the Genomic Era. Molecular Biology and Evolution 37:1530–1534.

Mo YK, Lanfear R, Hahn MW, Minh BQ. 2023. Updated site concordance factors minimize effects of homoplasy and taxon sampling. Bioinformatics 39:btac741.

Morrison DA, Morgan MJ, Kelchner SA. 2015. Molecular homology and multiple-sequence alignment: an analysis of concepts and practice. Australian Systematic Botany 28:46–62.

Mutti G, Ocaña-Pallarès E, Gabaldón T. 2025. Newly Developed Structure-Based Methods Do Not Outperform Standard Sequence-Based Methods for Large-Scale Phylogenomics. Mol Biol Evol 42.

Notredame C, Higgins DG, Heringa J. 2000. T-Coffee: A novel method for fast and accurate multiple sequence alignment. J Mol Biol 302:205–217.

Penn O, Privman E, Ashkenazy H, Landan G, Graur D, Pupko T. 2010. GUIDANCE: a web server for assessing alignment confidence scores. Nucleic Acids Res 38:W23–28.

Portik DM, Wiens JJ. 2021. Do Alignment and Trimming Methods Matter for Phylogenomic (UCE) Analyses? Systematic Biology 70:440–462.

Privman E, Penn O, Pupko T. 2012. Improving the performance of positive selection inference by filtering unreliable alignment regions. Mol Biol Evol 29:1–5.

Robinson DF, Foulds LR. 1981. Comparison of phylogenetic trees. Mathematical Biosciences 53:131–147.

Rokas A, Carroll SB. 2006. Bushes in the Tree of Life. PLOS Biology 4:e352.

Schneider A, Souvorov A, Sabath N, Landan G, Gonnet GH, Graur D. 2009. Estimates of Positive Darwinian Selection Are Inflated by Errors in Sequencing, Annotation, and Alignment. Genome Biology and Evolution 1:114–118.

Sela I, Ashkenazy H, Katoh K, Pupko T. 2015. GUIDANCE2: accurate detection of unreliable alignment regions accounting for the uncertainty of multiple parameters. Nucleic Acids Research 43:W7–W14.

Sharma S, Kumar S. 2022. Taming the Selection of Optimal Substitution Models in Phylogenomics by Site Subsampling and Upsampling. Molecular Biology and Evolution 39:msac236.

Shim H, Larget B. 2018. BayesCAT: Bayesian Co-estimation of Alignment and Tree. Biometrics 74:270–279.

Spielman SJ, Dawson ET, Wilke CO. 2014. Limited Utility of Residue Masking for Positive-Selection Inference. Molecular Biology and Evolution 31:2496–2500.

Spielman SJ, Miraglia ML. 2021. Relative model selection of evolutionary substitution models can be sensitive to multiple sequence alignment uncertainty. BMC Ecology and Evolution 21:214.

Stolzer M, Lai H, Xu M, Sathaye D, Vernot B, Durand D. 2012. Inferring duplications, losses, transfers and incomplete lineage sorting with nonbinary species trees. Bioinformatics 28:i409–i415.

Tan G, Muffato M, Ledergerber C, Herrero J, Goldman N, Gil M, Dessimoz C. 2015. Current Methods for Automated Filtering of Multiple Sequence Alignments Frequently Worsen Single-Gene Phylogenetic Inference. Syst Biol 64:778–791.

Thompson JD, Koehl P, Ripp R, Poch O. 2005. BAliBASE 3.0: latest developments of the multiple sequence alignment benchmark. Proteins 61:127–136.

Upham NS, Esselstyn JA, Jetz W. 2019. Inferring the mammal tree: Species-level sets of phylogenies for questions in ecology, evolution, and conservation. PLOS Biology 17:e3000494.

Wallace IM, O’Sullivan O, Higgins DG, Notredame C. 2006. M-Coffee: combining multiple sequence alignment methods with T-Coffee. Nucleic Acids Res 34:1692–1699.

Whelan S, Irisarri I, Burki F. 2018. PREQUAL: detecting non-homologous characters in sets of unaligned homologous sequences. Bioinformatics 34:3929–3930.

Wong KM, Suchard MA, Huelsenbeck JP. 2008. Alignment Uncertainty and Genomic Analysis. Science 319:473–476.

Wu M, Chatterji S, Eisen JA. 2012. Accounting for alignment uncertainty in phylogenomics. PLoS One 7:e30288.

Zhan Q, Ye Y, Lam T-W, Yiu S-M, Wang Y, Ting H-F. 2015. Improving multiple sequence alignment by using better guide trees. BMC Bioinformatics 16:S4.

Zhang C, Rabiee M, Sayyari E, Mirarab S. 2018. ASTRAL-III: polynomial time species tree reconstruction from partially resolved gene trees. BMC Bioinformatics 19:153.

Zhang C, Zhao Y, Braun EL, Mirarab S. 2021. TAPER: Pinpointing errors in multiple sequence alignments despite varying rates of evolution. Methods in Ecology and Evolution 12:2145–2158.

